# Evaluating the “cost of generating force” hypothesis across frequency in human running and hopping

**DOI:** 10.1101/2022.03.09.483693

**Authors:** Stephen P. Allen, Owen N. Beck, Alena M. Grabowski

## Abstract

The volume of active muscle and duration of extensor muscle force well-explain the associated metabolic energy expenditure across body mass and speed during level-ground running and hopping. However, if these parameters fundamentally drive metabolic energy expenditure, then they should pertain to multiple modes of locomotion and provide a simple framework for relating biomechanics to metabolic energy expenditure in bouncing gaits. Therefore, we evaluated the ability of the ‘cost of generating force’ hypothesis to link biomechanics and metabolic energy expenditure during human running and hopping across step frequencies. We asked participants to run and hop at 0%, ±8% and ±15% of preferred step frequency. We calculated changes in active muscle volume, force duration, and metabolic energy expenditure. Overall, as step frequency increased, active muscle volume decreased due to postural changes via effective mechanical advantage (EMA) or duty factor. Accounting for changes in EMA and muscle volume better related to metabolic energy expenditure during running and hopping at different step frequencies than assuming a constant EMA and muscle volume. Thus, to ultimately develop muscle mechanics models that can explain metabolic energy expenditure across different modes of locomotion, we suggest more precise measures of muscle force production that include the effects of EMA.

## Introduction

For decades, biomechanists and physiologists have sought to link the mechanics of running and hopping with the corresponding metabolic energy expenditure. One prevailing approach is the ‘cost of generating force’ hypothesis, which was proposed by Taylor and colleagues (Kram and Taylor, 1990; Taylor, 1994; Taylor et al., 1980) and posits that the primary determinant of metabolic energy expenditure required for running and hopping is the cost of generating muscle force to support body weight. This hypothesis is predicated on the fact that animals produce stride-average vertical ground reaction forces equal to body weight when running or hopping on level ground. Previous studies have demonstrated that metabolic energy expenditure depends on animal size, and that metabolic energy expenditure increases in almost direct proportion to the total weight of a running animal (Taylor et al., 1980). Further, per unit of body mass, it is more metabolically costly for smaller animals (e.g., mouse) to generate a unit of force than larger animals (e.g., horse) (Taylor, 1985), because small animals take more frequent strides and use less economical muscle fibers to produce force quickly (Heglund and Taylor, 1988). Thus, the metabolic energy expenditure during running and hopping varies with size and may depend on the number of strides taken per second, or stride frequency.

Kram and Taylor (Kram and Taylor, 1990) expanded the ‘cost of generating force’ hypothesis to explain why metabolic energy expenditure increases near linearly when running or hopping at faster velocities. They reasoned that the rate of force generation (i.e., the rate of cross bridge cycling) could be approximated by the inverse of ground contact time and formally proposed that the rate of metabolic energy expenditure (Ė_met_ in Watts) during running equals an animal’s body weight (*F*_*BW*_) multiplied by the inverse of ground contact time (*t*_*c*_^−1^) and a metabolic cost coefficient (*c*) (Eqn. 1).

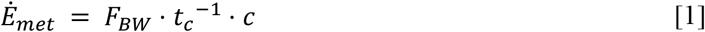

To produce the force needed to support body weight over each stride, animals need to activate a volume of muscle (i.e., the number of active actin-myosin crossbridges), which is primarily influenced by body weight and the leg’s effective mechanical advantage (EMA). EMA is the ratio of the ground reaction force moment arm to the muscle tendon moment arm. Kram and Taylor assumed that active muscle volume and EMA were independent of velocity (Biewener, 1989), which is why they simplified the equation to use force in units of body weight. Using this assumption, equation 1 well-described the increase in metabolic energy expenditure for a 10-fold increase in velocity and 4500-fold increase in body weight during forward hopping, trotting, and running animals (Kram and Taylor, 1990; Roberts et al., 1998a).

Since Kram and Taylor (Kram and Taylor, 1990), multiple studies have shown that active muscle volume and EMA change across running velocity and limb morphology (Kipp et al., 2018b; Roberts et al., 1998b; Wright and Weyand, 2001). Notably, Roberts et al. (Roberts et al., 1998b) demonstrated that running bipeds have a greater EMA than size-matched quadrupeds due to their upright posture, which influences active muscle volume and metabolic energy expenditure. Thus, the authors proposed a refined version of the ‘cost of generating force’ hypothesis to account for changes in active muscle volume where the rate of metabolic energy expenditure equals the product of active muscle volume (*V*_*m*_), the inverse of ground contact time, and a new cost coefficient (*k*) (Eqn. 2)

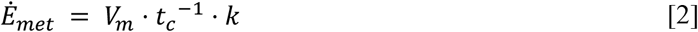

Kipp et al. (Kipp et al., 2018b) applied this refined version of the ‘cost of generating force’ hypothesis (Eqn. 2) to human running and found that humans decrease their EMA and increase active muscle volume by as much as 53% from 2.2 m·s^-1^ to 5.0 m·s^-1^. Thus, the authors concluded that the curvilinear increase in metabolic energy expenditure with running velocity (Batliner et al., 2018) results from an increase in active muscle volume and an increase in the rate of force production due to shorter ground contact times.

Though the rate of force generation and active muscle volume well-explain metabolic energy expenditure across different running and hopping velocities, it is unknown if these biomechanical variables explain changes in metabolic energy expenditure across different stride and step frequencies, where a step equals ground contact and the subsequent aerial time and two steps comprise a stride. Previous studies have shown that humans have a preferred step frequency for running and hopping that minimizes metabolic energy expenditure, and deviating from the preferred step frequency increases metabolic energy expenditure (Allen and Grabowski, 2019; Cavagna et al., 1988; Cavanagh and Williams, 1982; Farris and Sawicki, 2012; Grabowski and Herr, 2009; Högberg, 1952; Raburn et al., 2011; Swinnen et al., 2021); thus there is a U-shaped relationship between metabolic energy expenditure and step frequency (Doke and Kuo, 2007; Snyder and Farley, 2011; Swinnen et al., 2021). When considering the ‘cost of generating force’ hypothesis, Gutmann and Bertram (Gutmann and Bertram, 2017a; Gutmann and Bertram, 2017b) suggest that the rate of force production alone (Eqn. 1) cannot fully account for the U-shaped changes in metabolic energy expenditure with hopping frequency. Instead, accounting for changes in active muscle volume and the rate of force production (Eqn. 2) may better explain this U-shaped relationship. Previous studies have suggested that the U-shaped relationship is due to simultaneous increasing and decreasing metabolic costs (Doke and Kuo, 2007; Snyder and Farley, 2011; Swinnen et al., 2021) where ground contact time decreases with increased step frequency during human running and hopping, which implies that humans must produce forces at a faster rate and increase metabolic cost (Farley et al., 1991). Simultaneously, increased step frequencies are accompanied by shorter steps during running and less center of mass displacement during running and hopping, both of which may increase EMA and reduce active muscle volume and decrease metabolic cost (Monte et al., 2021). Thus, accounting for changes in the rate of force production and active muscle volume through EMA may better describe metabolic energy expenditure across step frequencies than the cost of generating force alone. We hypothesized that accounting for changes in active muscle volume and the rate of force production (Eqn. 2) better explains changes in metabolic energy expenditure across step frequencies compared to the original “cost of generating force” equation, which estimates active muscle volume from body weight (Eqn. 1) for both running and hopping. Further, we hypothesize that active muscle volume decreases as step frequency increases in running and hopping due to increased EMA.

## Materials and Methods

### 1. Participants

Ten healthy runners (6F, 4M; average ± s.d., mass: 60.7 ± 8.9 kg, height: 1.72 ± 0.09 m, age: 24.5 ± 3.4 years) with no reported cardiovascular, neurological, or musculoskeletal impairments participated in the study. All participants reported running for exercise at least 30 minutes per day, three times per week, for at least 6 months. Each participant provided written informed consent to participate in the study according to the University of Colorado Boulder Institutional Review Board.

### 2. Experimental Protocol

Over two separate days, participants performed a series of running trials on a force-measuring treadmill (Treadmetrix, Park City, UT; 1000 Hz) and stationary, two-legged hopping trials on force plates (Bertec, Columbus, OH; 1000 Hz) while we simultaneously measured ground reaction forces, lower limb kinematics, and metabolic energy expenditure throughout each trial. On the first day, participants performed six 5-min running trials at 3 m·s^-1^. During the first trial, we determined each participant’s preferred step frequency (PSF). We collected ground reaction forces (GRFs) for 15-sec during the third and fifth minute of the first trial and determined average PSF from ground contact events identified by a 20 N vertical GRF threshold. We then instructed participants to complete the remaining running trials while matching their step frequency to the timing of an audible metronome. The metronome was set to 85%, 92% 100%, 108% and 115% of their PSF, similar to previous studies (Snyder and Farley, 2011; Swinnen et al., 2021), and the order of the trials was randomized.

On the second day, participants performed five, 5-min stationary hopping trials, on both feet. To account for the effects of frequency on metabolic energy expenditure and given the similarity of frequencies that minimize metabolic energy expenditure during hopping and running (Allen and Grabowski, 2019; Cavagna et al., 1997; Farris and Sawicki, 2012; Grabowski and Herr, 2009; Kaneko et al., 1987), we instructed participants to hop in place while matching their step frequency to the audible metronome set to 85%, 92% 100%, 108% and 115% of their PSF from day 1. The order of the hopping trials was randomized, and we did not determine preferred hopping frequency.

### 3. Metabolic Energy Expenditure

We measured participants’ rates of oxygen consumption and carbon dioxide production via indirect calorimetry (TrueOne 2400, ParvoMedics, Sandy, UT) throughout each running and hopping trial. We instructed participants to refrain from exercising before each experimental session or ingesting caffeine four hours before each experimental session to minimize day-to-day variability in metabolic rates. Additionally, participants were instructed to be at least two hours postprandial at the start of each experimental session to mitigate potential effects of diet on metabolic measurements. Further, each experimental session was performed at the same time each day and separated by at least 24 hours to eliminate any potential effects of day-to-day variability or fatigue. We calculated gross steady-state metabolic power from the average metabolic rates during the last two minutes of each 5-min trial using a standard equation (Kipp et al., 2018a; Péronnet and Massicotte, 1991).

### 4. Kinematic and Kinetics

We positioned 40 reflective markers bilaterally on both legs and the pelvis. Markers on the ankles and knees were used to define joint centers and clusters of 3-4 markers were placed on each segment prior to experimental trials. We collected lower limb kinematic data for 15-sec during the last minute of each trial using 3-dimensional motion capture (Vicon Nexus 2.3, Oxford, UK; 200 Hz) simultaneously with GRFs. We analyzed 20 steps from each trial and used a 4^th^ order low-pass Butterworth filter with a 20 Hz cut-off to process analog GRF signals and marker trajectories (Alcantara, 2019; Mai and Willwacher, 2019). We determined ground contact using a 20 N vertical GRF threshold for both running and hopping and calculated the rate of force production as the inverse of ground contact time (*t*_*c*_^*-1*^).

To calculate EMA and *V*_*m*_, we estimated the average extensor muscle-tendon unit force (*F*_*mtu*_) about each joint assuming a constant muscle-tendon moment arm (*r*) for each muscle group and using instantaneous ankle, knee, and hip sagittal joint moments from Visual 3D (Visual 3D, C-Motion Inc., Germantown, MD, USA) (Biewener et al., 2004; Kipp et al., 2018b). We only included joint moment values that exceeded 25% of the maximum extensor moment due to the inherently noisy center of pressure measurements caused by low force values at the beginning and end of the ground contact phase (Biewener et al., 2004; Griffin et al., 2003; Kipp et al., 2018b). Because the net joint moments of the knee and hip include flexion moments from bi-articular muscles, we accounted for forces in bi-articular muscles by assuming *F*_*mtu*_ was proportional to physiological cross-sectional area of active muscle fibers (Eq. 3-5).

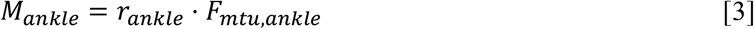

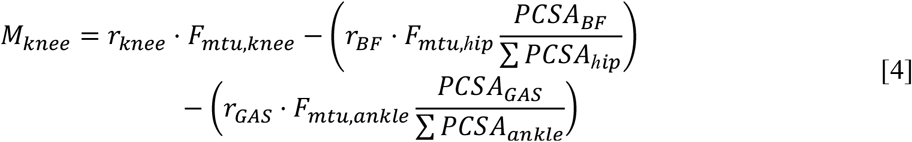

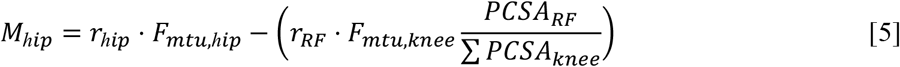

where *M* is the net joint moment, *r* is a weighted-average muscle-tendon moment arm, and *PCSA* is the physiological cross-sectional area. *GAS, BF*, and *RF* represent the properties of the gastrocnemius, biceps femoris, and rectus femoris muscles, respectively. We calculated *F*_*mtu,ankle*_ from Eqn. 3, and solved Eqn. 4 and 5 simultaneously due to the two unknown quantities of *F*_*mtu*.*knee*_ and *F*_*mtu,hip*_. We considered moments that extend joints to be positive. Values for *r* and *PCSA* were taken from the anthropometric data of four male human cadavers reported in Biewener et al. (Biewener et al., 2004) and previously used in Kipp et al. (Kipp et al., 2018b). We then used the quotient of the average sagittal plane resultant GRF magnitude and *F*_*mtu*_ at each joint during ground contact to calculate EMA, which equals the quotient of *r* and the GRF moment arm (*R*).

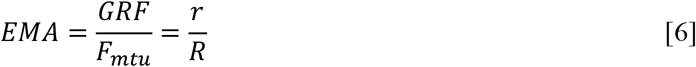

We calculated *V*_*m*_ separately for each joint (Eqn. 7) and then summed them to estimate the total average *V*_*m*_ per leg. To do this, we assumed the muscles produced force isometrically with a constant stress (σ = 20 N·cm^-2^) (Perry et al., 1988) and combined this with our estimates of *F*_*mtu*_ and weighted-average fascicle length (*L*) from Biewener et al. (Biewener et al., 2004),

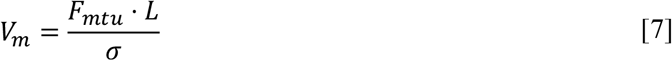

### 5. Estimating cost-coefficients and metabolic energy expenditure

We calculated the metabolic cost-coefficients, *c* and *k*, for each trial during running and hopping using Eqns. 1 and 2. We averaged each cost-coefficient across the range of frequencies (separately for running and hopping). Then we implemented the respective cost-coefficient averages in addition to *V*_*m*_, *t*_*c*_^*-1*^, and *F*_*BW*_ to predict metabolic power for each step frequency during running and hopping using Eqns. 1 and 2.

### 6. Statistics

To evaluate the agreement between measured metabolic power and predicted metabolic power from Eqns. 1 & 2, we performed limits of agreement analyses (Bland-Altman) for each target step frequency and calculated the systematic bias (mean differences) and 95% limits of agreement. We also constructed linear mixed-effects models (α = 0.05) to determine the effect of measured step frequency relative to PSF on *t*_*c*_^*-1*^, *c, k, EMA, V*_*m*_, average joint extensor moment, and average sagittal plane resultant GRF magnitude. In each linear mixed-effects model, we considered measured step frequency relative to PSF as a fixed effect and participant as a random effect. Model coefficients are reported alongside their p-values and represent the change in the dependent variable per a 1% change in measured step frequency relative to PSF. We performed all statistical analyses in R (version 3.6.3) (R Core Team, 2020) using custom scripts and packages (Datta, 2017; Pinheiro et al., 2020; Revelle, 2019; Wickham, 2016).

## Results

We removed data for one participant at the 85% PSF and 92% PSF running trials because they were >3% off of the target step frequencies.

### 1. Running

On average, measured metabolic power was minimized when participants ran at their PSF (Fig. 1A), which was a step frequency of (avg. ± s.d.) 2.90 ± 0.09 Hz (Table 1). As participants deviated from their PSF, average measured metabolic power increased by 17% and 9% when running at 85% of PSF and 115% of PSF, respectively (Fig. 1A). Overall, metabolic power estimated with Eqn. 1 underestimated average metabolic power for step frequencies slower than PSF (up to 13% at 85% PSF) but overestimated average metabolic power for step frequencies equal to or greater than PSF (up to 9.5% greater at 108% PSF) (Fig. 2A & 3A). Limits of agreement analysis show metabolic power estimated with Eqn. 2 had a bias closer to zero and lower than Eqn. 1 at each step frequency, however, the magnitude of the upper and lower limits of agreement for Eqn. 2 were greater than those of Eqn. 1 due to increased variability (Fig. 2A & 3A) during running.

**Figure 1:**
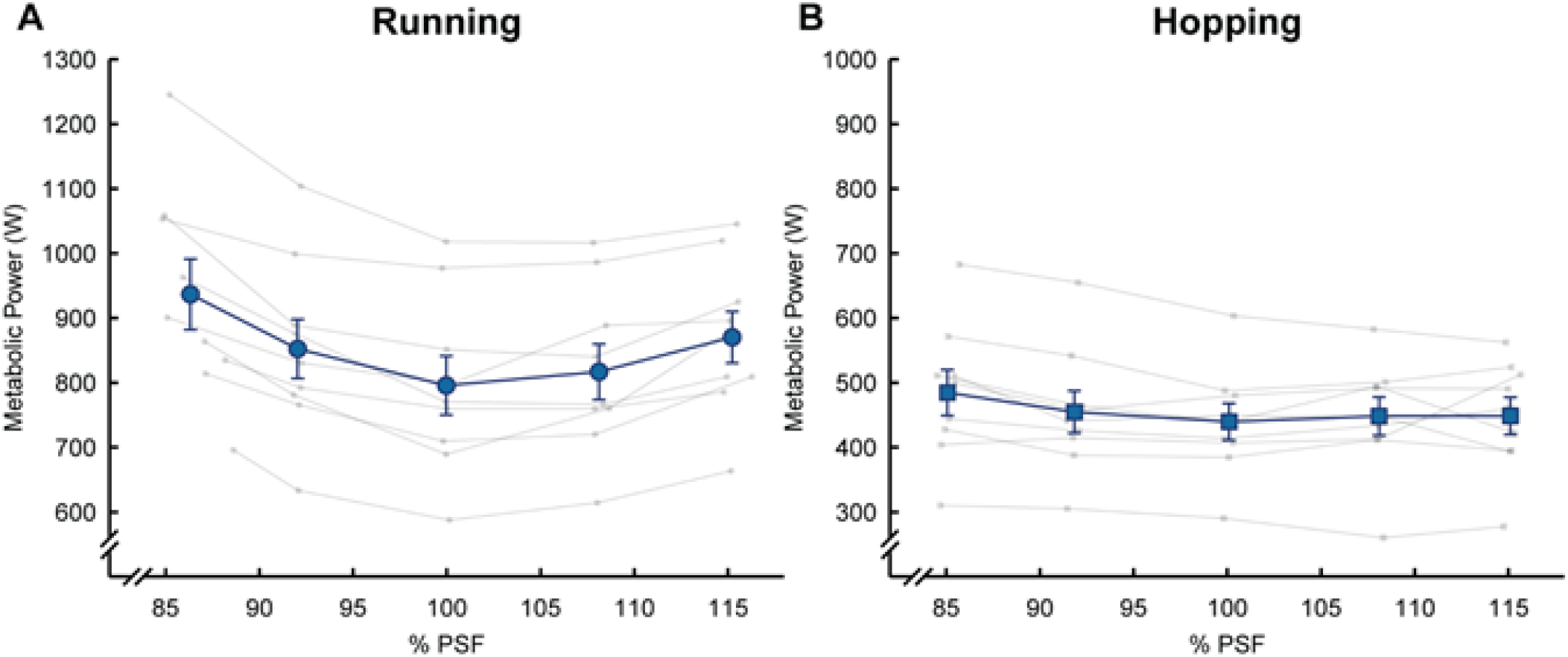
Gross metabolic power across percentage of preferred step frequency. Average ± s.e.m. power (metabolic large, blue symbols) and values from individual subjects (small, grey symbols) versus the percentage of running preferred step frequency (% PSF) in A) running and B) hopping. Vertical and horizontal error bars may not be visible behind data points.

**Table 1.** Biomechanical variables for running (3 m□s^-1^) and hopping in place at different percentages of preferred running step frequency.

|  | Target<br>% PSF | Achieved<br>% PSF | Achieved step<br>frequency<br>(Hz) | Stance Avg.<br>resultant GRF<br>(N) | Avg. extension moment (N·m) |  |  |
| --- | --- | --- | --- | --- | --- | --- | --- |
|  |  |  |  |  | Ankle | Knee | Hip |
| Running | 85 | $86.3 \pm 1.5$ | $2.50 \pm 0.1$ | $939 \pm 165$ | $142 \pm 32$ | $106 \pm 30$ | $64 \pm 14$ |
| | 92 | $92.0 \pm 0.2$ | $2.67 \pm 0.1$ | $890 \pm 154$ | $134 \pm 33$ | $103 \pm 26$ | $58 \pm 12$ |
| | 100 | $100.0 \pm 0.1$ | $2.90 \pm 0.1$ | $848 \pm 137$ | $127 \pm 27$ | $95 \pm 23$ | $52 \pm 12$ |
| | 108 | $108.1 \pm 0.3$ | $3.13 \pm 0.1$ | $831 \pm 117$ | $123 \pm 25$ | $81 \pm 24$ | $52 \pm 14$ |
| | 115 | $115.2 \pm 0.5$ | $3.34 \pm 0.1$ | $807 \pm 102$ | $122 \pm 17$ | $70 \pm 20$ | $54 \pm 16$ |
| Hopping | 85 | $85.1 \pm 0.4$ | $2.46 \pm 0.1$ | $889 \pm 134$ | $128 \pm 30$ | $118 \pm 28$ | $37 \pm 15$ |
| | 92 | $91.8 \pm 0.3$ | $2.66 \pm 0.1$ | $877 \pm 139$ | $130 \pm 39$ | $102 \pm 27$ | $35 \pm 12$ |
| | 100 | $100.1 \pm 0.2$ | $2.90 \pm 0.1$ | $883 \pm 121$ | $137 \pm 33$ | $75 \pm 24$ | $34 \pm 11$ |
| | 108 | $108.1 \pm 0.2$ | $3.13 \pm 0.1$ | $870 \pm 127$ | $130 \pm 33$ | $57 \pm 18$ | $37 \pm 12$ |
| | 115 | $115.1 \pm 0.3$ | $3.33 \pm 0.1$ | $868 \pm 131$ | $132 \pm 30$ | $41 \pm 16$ | $37 \pm 13$ |

Average ± s.d.; PSF, preferred running step frequency; GRF, ground reaction force; Avg. joint moment is defined when extension moments are greater than 25% of the peak joint moment.

**Figure 2:**
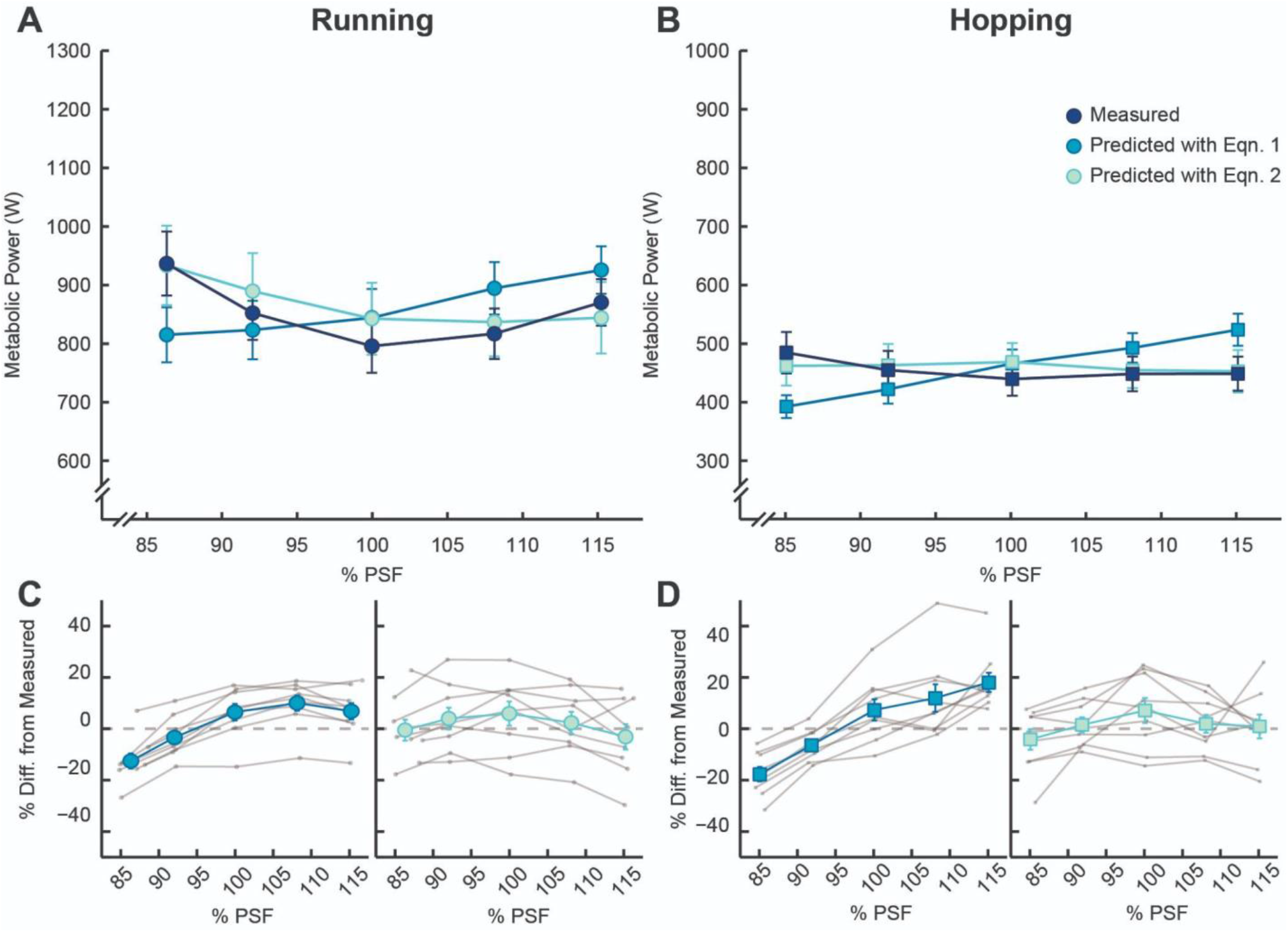
Predicted gross metabolic power across percentage of preferred step frequency. Average ± s.e.m. (large, colored symbols) gross metabolic power for measured (dark blue) and predicted values using Eq.1 (blue) and Eqn. 2 (light blue) versus the percentage of running preferred step frequency (% PSF) in A) running and B) hopping. Vertical and horizontal error bars may not be visible behind the data points. C) Running and D) hopping percent difference between each equation and measured metabolic power for average ± s.e.m. and participants (small, grey symbols).

**Figure 3:**
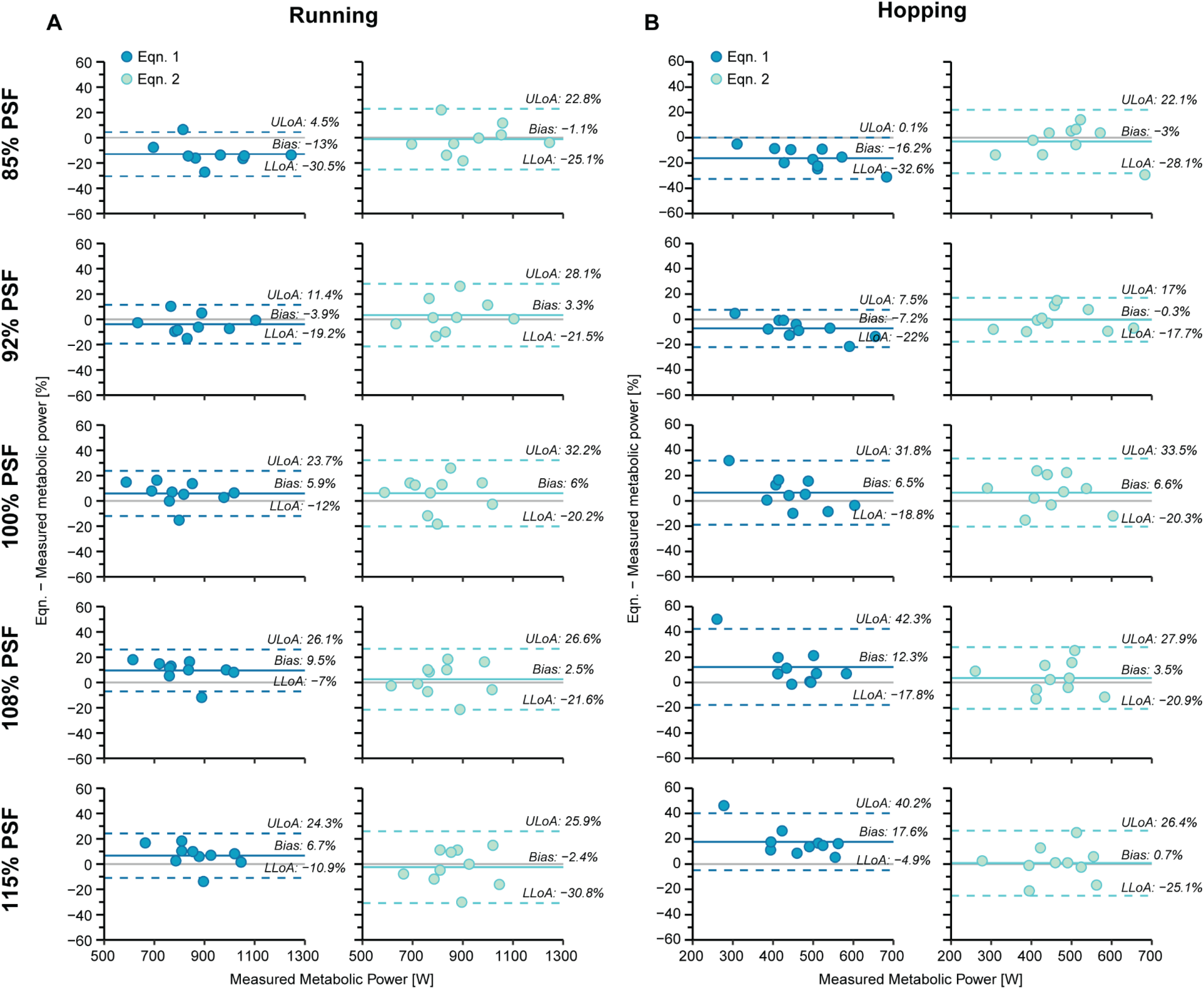
Limits of Agreement (Bland-Altman). Limits of agreement analysis comparing the percent difference between Eqn. 1 (blue symbols) or Eqn. 2 (light blue symbols) and gross metabolic power measured via indirect calorimetry. Mean differences (Bias) are indicated by the solid-colored lines, while the lower and upper limits of agreement (LLoA/ULoA) are denoted by dashed-colored lines. LLoA/ULoA were calculated using 1.96 SD.

The linear mixed-effects model showed that total *V*_*m*_ decreased by 20.54 cm^3^ for every 1% increase in step frequency relative to PSF (p<0.001; Fig. 4; Table 2). Specifically, participants decreased ankle, knee, and hip *V*_*m*_ by 3.68 cm^3^, 10.34 cm^3^, and 5.54 cm^3^, respectively, for every 1% increase in step frequency (p<0.001 for each; Fig. 4; Table. 2). Despite the reduction in joint-specific *V*_*m*_, we did not detect significant changes in ankle, knee, or hip EMA across step frequency (p=0.66; p=0.05; p=0.59, respectively). Average (± s.d.) EMA across step frequencies for the ankle, knee, and hip was 0.314 ± 0.017, 0.393 ± 0.084, and 0.714 ± 0.117, respectively (Fig. 5, Table 2). Rather, the changes in joint-specific *V*_*m*_ may have been due to the decrease in average ankle, knee, and hip extensor moments as step frequency increased. Average ankle, knee, and hip extensor moments decreased by 0.66 N·m (p<0.001), 1.3 N·m (p<0.001), and 0.31 N·m (p<0.01), respectively, for every 1% increase in step frequency (Table 1). Finally, *t*_*c*_^*-1*^ increased by 0.02 s^-1^ for every 1% increase in step frequency relative to PSF during running (p<0.001; Fig. 6A). We used these variables to solve for the cost-coefficient and found that *c* decreased by 0.003 J·N^-1^ for every 1% increase in step frequency (p<0.001; Fig. 7A), but *k* did not change across step frequency, and averaged (± s.d.) 0.087 ± 0.003 J·cm^-3^ (p=0.18; Fig. 7A).

**Figure 4:**
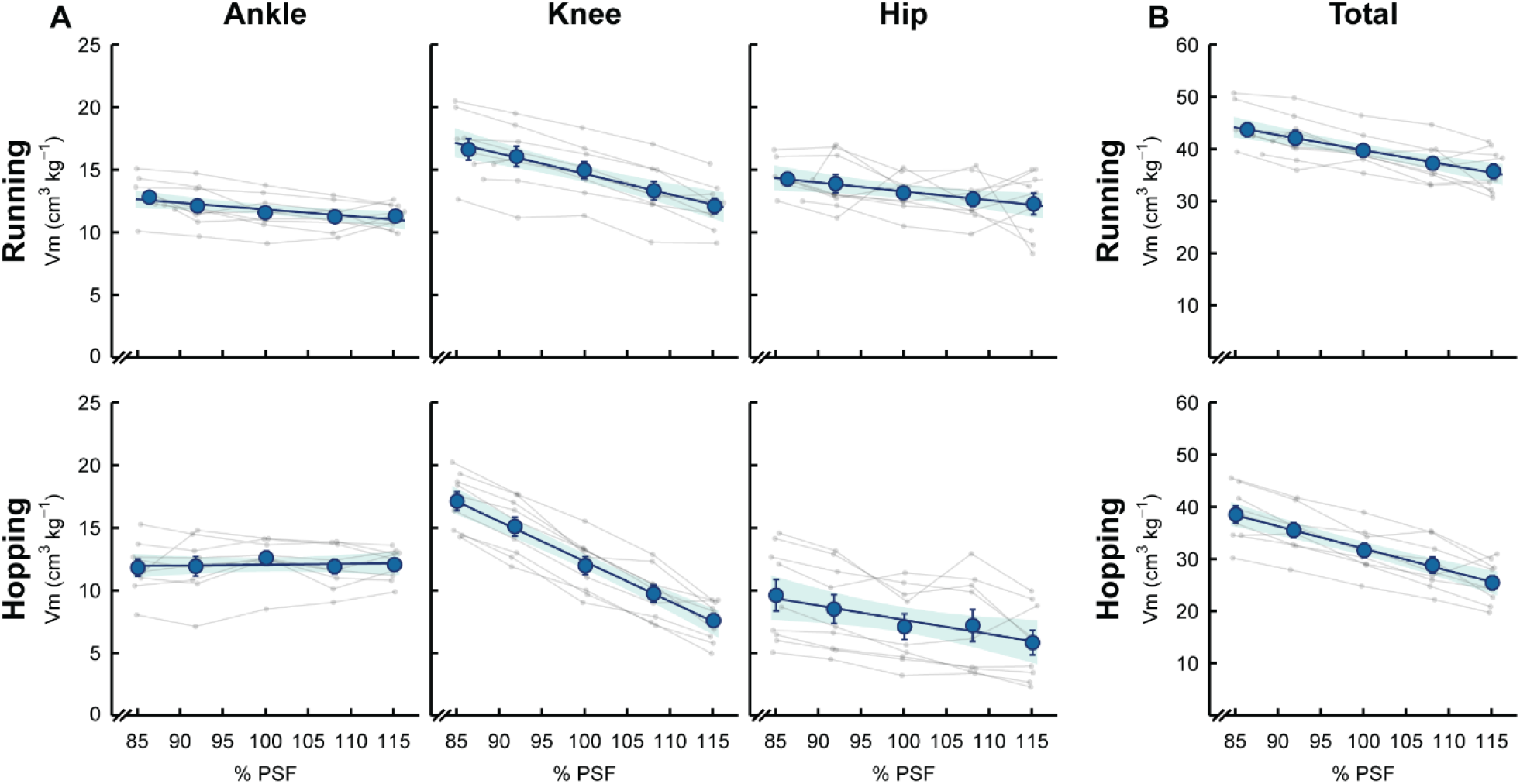
Active muscle volume across percentage of preferred step frequency. Average ± s.e.m. active muscle volume (*Vm*) of the leg extensors during ground contact (large, blue symbols) and values from individual subjects (small, grey symbols) versus the percentage of preferred running step frequency (% PSF) for running and hopping. A) *Vm* of the muscles surrounding the ankle, knee, and hip joints during running and hopping, and B) is the summed total of the ankle, knee, and hip joint *Vm*. The dark lines represent the results of linear mixed-effects models, and the shaded regions represent the model’s 95% confidence intervals. Coefficients and intercepts for each of the linear mixed-effects models are presented in Table 2. Vertical and horizontal error bars may not be visible behind data points.

**Table 2.** Linear mixed-effects model results for effective mechanical advantage (Fig. 5) and active muscle volume (Fig. 4) at the ankle, knee, hip, and summed total while running and hopping at different percentages of preferred running step frequency.

| | | EMA | | | $V_m$ | | |
| --- | --- | --- | --- | --- | --- | --- | --- |
|  |  | Intercept | Slope | p-value | Intercept | Slope | p-value |
| Running | Ankle | 0.32 | $-0.24 \cdot 10^{-5}$ | 0.87 | 1095.6 | -3.8 | <0.001 |
| | Knee | 0.29 | $1.0 \cdot 10^{-3}$ | 0.07 | 1896.3 | -10.2 | <0.001 |
| | Hip | 0.76 | $-4.9 \cdot 10^{-4}$ | 0.67 | 1354.6 | -5.8 | <0.001 |
|  | Total | - | - | - | 4345.2 | -19.7 | <0.001 |
| Hopping | Ankle | 0.39 | $-3.6 \cdot 10^{-4}$ | 0.06 | 655.1 | 0.7 | 0.32 |
| | Knee | -0.40 | $8.4 \cdot 10^{-3}$ | <0.001 | 2565.3 | -18.0 | <0.001 |
| | Hip | 0.63 | $2.7 \cdot 10^{-3}$ | 0.35 | 917.8 | -3.8 | <0.001 |
|  | Total | - | - | - | 4138.8 | -21.0 | 0.008 |

EMA: effective mechanical advantage; $V_m$ : active muscle volume; p-value < 0.05 indicates a slope significantly different from zero with respect to normalized stride frequency. Linear mixed-effects model is in the form of $Y = \text{Intercept} + \%PSF \cdot \text{Slope}$ .

**Figure 5:**
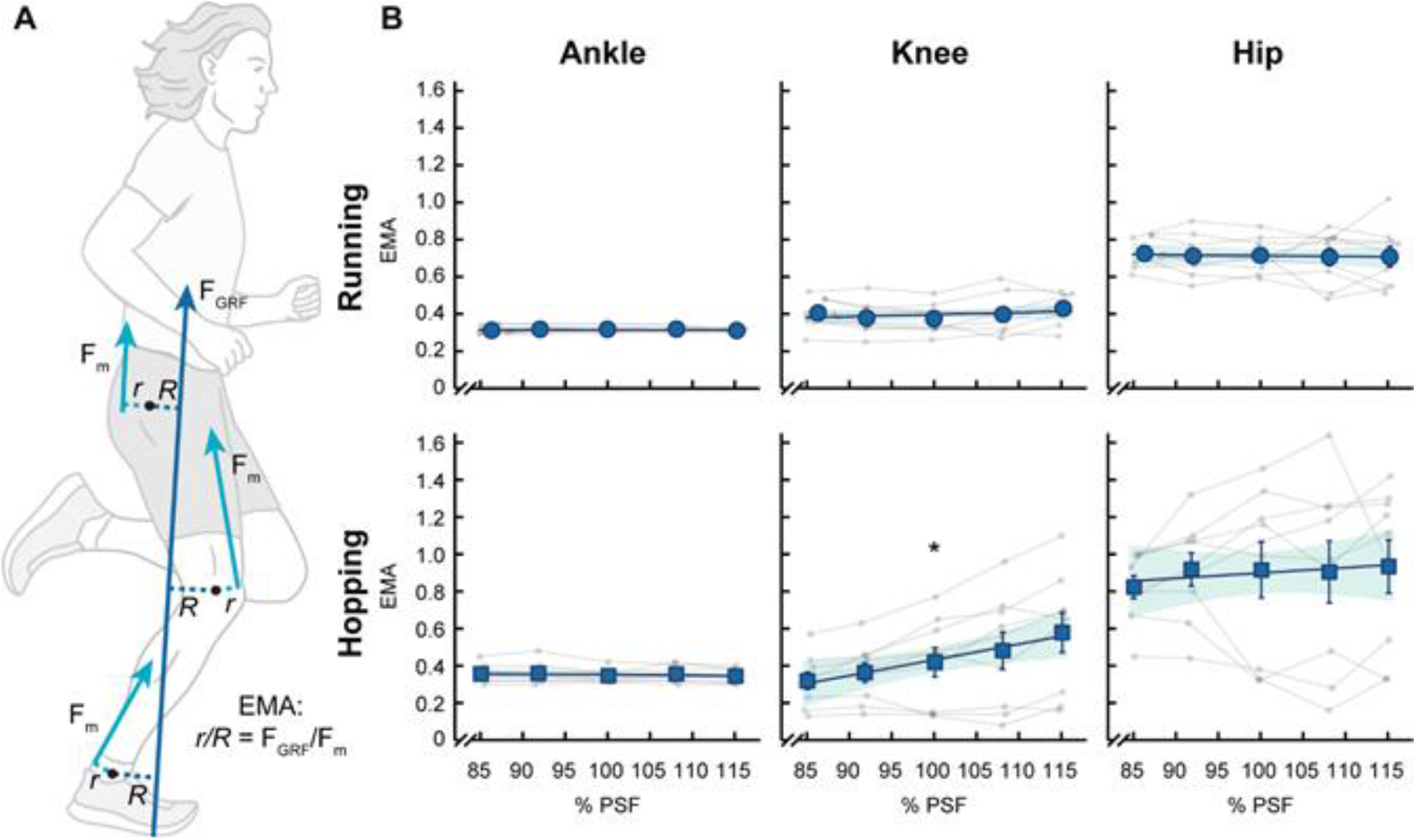
Effective mechanical advantage (EMA) across percentage of preferred step frequency. A) Illustration of EMA during running, which equals the ratio of the muscle-tendon moment arm (r) and the external resultant ground reaction force moment arm (R) or the ratio of resultant ground reaction force (FGRF) and muscle force (Fm). B) Average ± s.e.m. EMA for the ankle, knee, and hip joints (large, blue symbols) with values for individual subjects (small, grey symbols) versus the percentage of preferred running step frequency (% PSF). The dark lines represent the results of linear mixed-effects models and the shaded regions represent the model’s 95% confidence intervals. Coefficients and intercepts for each of the linear mixed-effects models are presented in Table 2. * indicates if the model slope is significantly different from zero. Vertical and horizontal error bars may not be visible behind data points.

**Figure 6:**
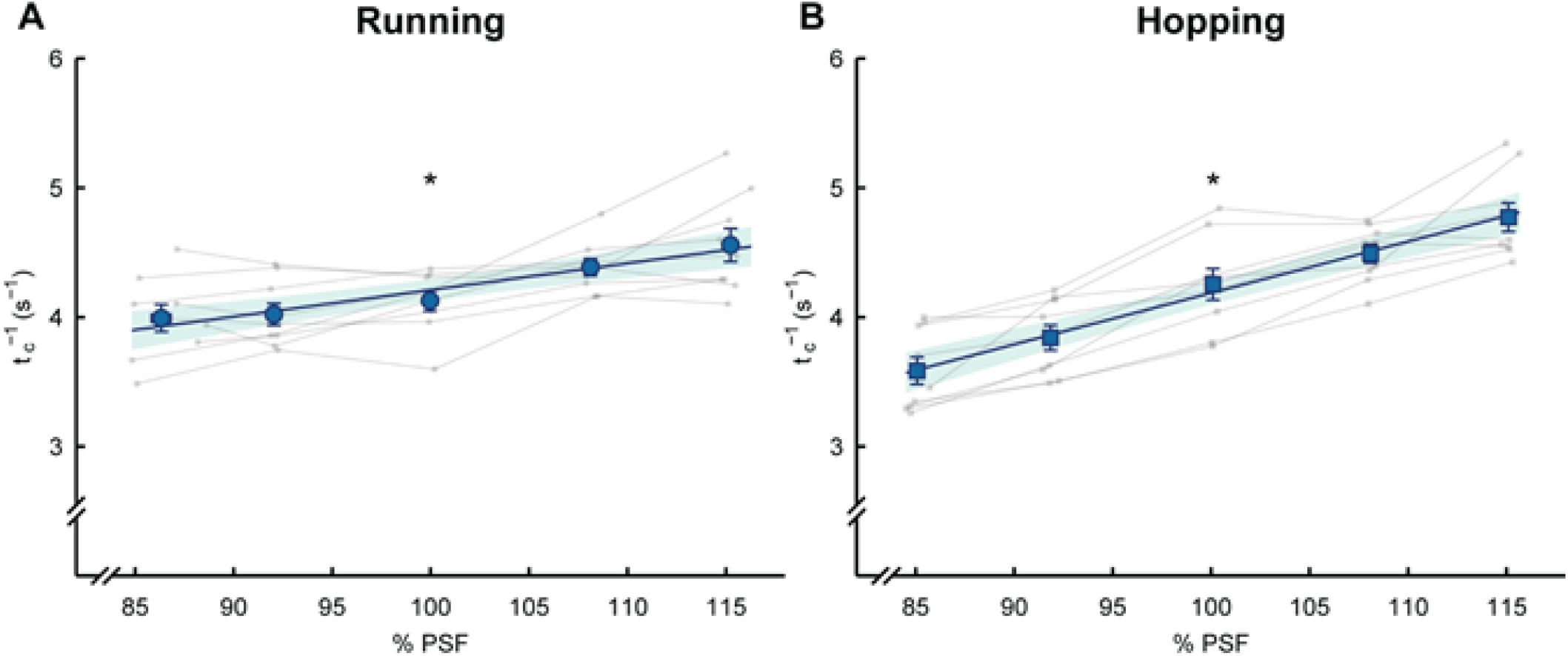
Rate of force production across percentage of preferred step frequency. Average ± s.e.m. rate of force production (*tc*-1; large, blue symbols) and values from individual subjects (small, grey symbols) versus the percentage of running preferred step frequency (% PSF) for A) running and B) hopping. The dark lines represent the model prediction across percentage of preferred step frequency (running: *tc*-1 = 0.021 · PSF + 2.160, hopping: *tc*-1 = 0.040 · PSF + 0.216) and the shaded areas represent the 95% confidence interval. * indicates if the model slope is significantly different from zero. Vertical and horizontal error bars may not be visible behind data points.

**Figure 7:**
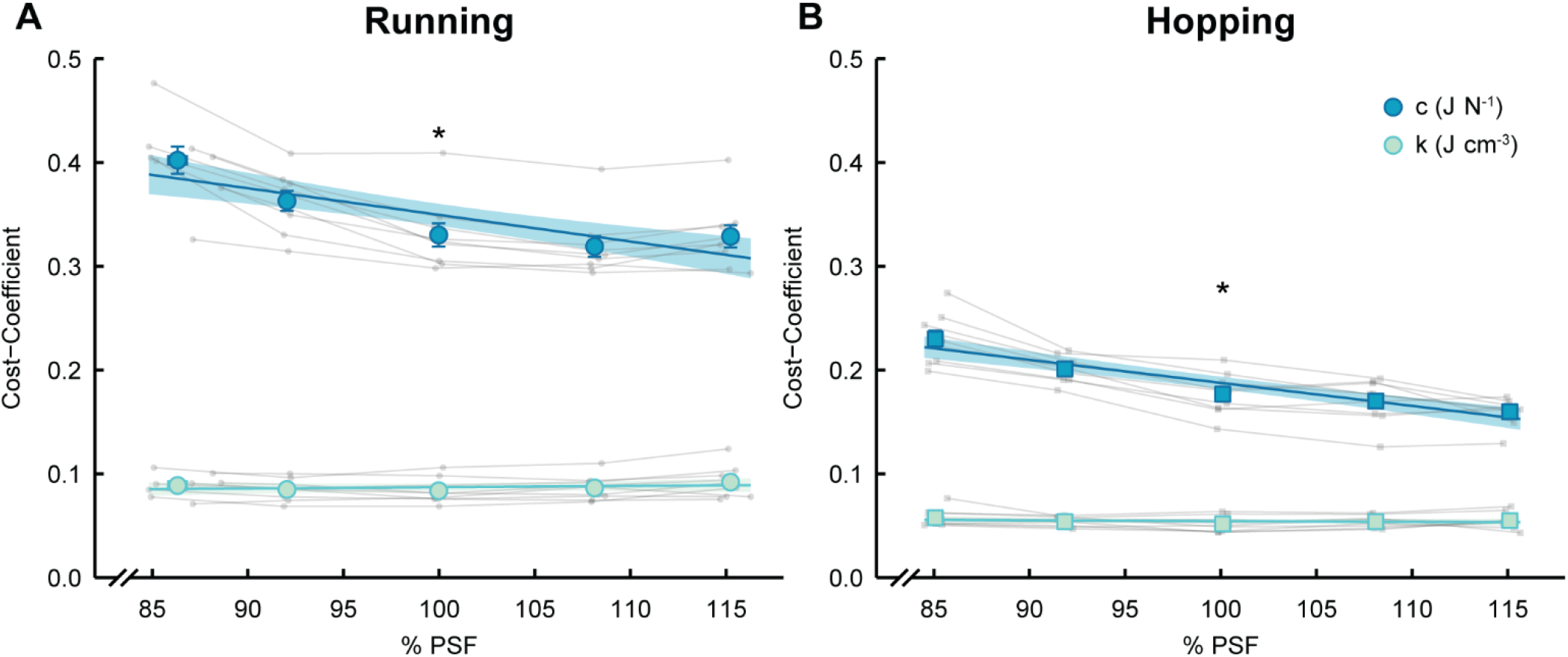
Cost-coefficient across percentage of preferred step frequency. Average ± s.e.m. cost coefficients (*c* – large, dark blue symbols and *k* – large, light blue symbols) and values from individual subjects (small, grey symbols) versus the percentage of running preferred step frequency (% PSF) in A) running and B) hopping. The lines represent the results of the linear mixed-effects model where *c* = -0.003 · PSF + 0.601 (p<0.001) and *k* = 1.16×10^−4^ · PSF + 0.075 (p=0.18) for running, and *c* = -0.0022 · PSF + 0.407 (p<0.001) and *k* = 0.001 · PSF + 0.045 (p=0.20) for hopping. The p-values indicate if the slope is significantly different than zero. The dark lines represent the results of linear mixed-effects models, and the shaded regions represent the model’s 95% confidence intervals. Vertical and horizontal error bars may not be visible behind data points.

### 2. Hopping

On average, measured metabolic power numerically increased by 10% and 2% when hopping at 85% of PSF and 115% of PSF, respectively, relative to 100% PSF (p>0.9; Fig. 1B). On average, metabolic power estimated with Eqn. 1 underestimated metabolic power for step frequencies slower than PSF (up to 16% at 85% PSF) but overestimated metabolic power for step frequencies greater than PSF (up to 17% at 115% PSF) (Fig. 2B & 3B). Metabolic power estimated with Eqn. 2 had a bias closer to zero and lower than Eqn. 1 at each step frequency (Fig. 2B & 3B). The magnitude of the upper and lower limits of agreement for Eqn. 2 were greater than those of Eqn. 1 due to increased variability during hopping (Fig. 2B & 3B).

The linear mixed-effects models showed that total *V*_*m*_ decreased by 21 cm^3^ for every 1% increase in step frequency relative to PSF (p<0.001; Fig. 4; Table 2). Participants decreased joint-specific *V*_*m*_ at the knee and hip by 18.0 cm^3^ and 3.7 cm^3^ for every 1% increase in step frequency (p<0.001 and p=0.008, respectively; Fig. 4), whereas ankle *V*_*m*_ did not change across step frequency and averaged (± s.d.) 729 ± 176 cm^3^ (p=0.32; Fig. 4). We found that knee EMA increased by 0.008 for every 1% increase in step frequency (p<0.001; Fig. 5, Table 2). However, ankle and hip EMA did not change across step frequency and averaged (± s.d.) 0.35 ± 0.04 (p=0.06) and 0.90 ± 0.37 (p=0.35), respectively (Fig. 5; Table 2). Similarly, participants decreased average knee extensor moment by 2.6 N·m for every 1% increase in step frequency (p<0.001; Table 1). However, average ankle and hip extensor moments did not change across step frequency and averaged (± s.d.) 129 ± 27 N·m (p=0.32) and 56 ± 13.9 N·m (p=0.77), respectively. Finally, *t*_*c*_^*-1*^ increased by 0.04 s^-1^ for every 1% increase in step frequency relative to PSF during stationary hopping (p<0.001; Fig 6B). We used these variables to solve for the cost-coefficients and found that *c* decreased by 0.0022 J·N^-1^ for every 1% increase in step frequency (p<0.001; Fig. 7B), but *k* did not change across step frequency and averaged (± s.d.) 0.054 ± 0.002 J·cm^-3^ (p=0.20; Fig. 7B).

## Discussion

Our data support our first hypothesis that accounting for changes in active muscle volume (*V*_*m*_) and the rate of force production (*t*_*c*_^*-1*^) (Eqn. 2) better explain changes in metabolic energy expenditure across step frequencies compared to the original “cost of generating force” equation, which estimates *V*_*m*_ through body weight (Eqn. 1). We also found that accounting for changes in *V*_*m*_ and *t*_*c*_^*-1*^ (Eqn. 2) results in a constant cost-coefficient, *k* (Fig. 7), across step frequencies for running and hopping. The average values for *k* (0.087 J□cm^-3^ and 0.056 J□cm^-3^ for running and hopping, respectively) are in line with previous values reported for human running (0.079 J□cm^-3^) at different velocities (Kipp et al., 2018b). Our data also support our second hypothesis, that *V*_*m*_ is reduced as step frequency increases in human running and stationary hopping. When step frequency increased from 85% PSF to 115% PSF, we found that *V*_*m*_ decreased by 18% and 26% during running and hopping, respectively. This reduction predominantly occurred due to changes at the knee in both running and hopping, with smaller or non-significant contributions from the ankle and hip during both tasks (Fig. 4; Table 2). We found that the knee accounted for ∼55% and ∼87% of the change in total *V*_*m*_ during running and hopping, respectively, whereas, when humans run at faster velocities from 2.2 – 5.0 m□s^-1^, the ankle, knee, and hip account for ∼39%, ∼20%, and ∼41% of the change in total active muscle volume, respectively (Kipp et al., 2018b). Our data, along with previous studies, support the general hypothesis that the metabolic cost of bouncing gaits is related to *V*_*m*_ recruited to generate force and the rate that the force is produced (Heglund and Taylor, 1988; Kipp et al., 2018b; Roberts et al., 1998b; Taylor et al., 1980; Wright and Weyand, 2001).

The mechanism by which total *V*_*m*_ decreased with step frequency differed between running and hopping. We found that joint-specific effective mechanical advantage (EMA) was independent of step frequency during running (Fig. 4; Table 2). Therefore, the reductions in total *V*_*m*_ during running were likely due to greater duty factors, which resulted in reduced stance-average resultant ground reaction forces (GRFs) and the corresponding joint moments (Table 1). In comparison, during hopping, EMA at the knee increased by 82% when step frequency increased from 85% to 115% PSF, while the magnitude of stance-average resultant GRF did not change (Fig. 5; Table 1 & 2). This might imply that participants decreased total *V*_*m*_ during hopping by altering their lower limb position to hop with a straighter leg and extended knee as step frequency increased. When taken together, these results suggest that humans may utilize two different mechanisms to alter total *V*_*m*_ during bouncing gaits, duty factor (Beck et al., 2020) and EMA. Previously, Kipp et al. (Kipp et al., 2018b) demonstrated that humans utilize both mechanisms simultaneously to increase total *V*_*m*_ when running at different velocities. They found that runners increased total *V*_*m*_ by 53% with faster running velocities from 2.2 – 5.0 m□s^-1^ due to a concurrent decrease in duty factor and decrease in hip EMA, which is likely due to the increased step frequency that accompanies faster running velocity (Heglund and Taylor, 1988).

Our measures of knee and ankle EMA during two-legged, stationary hopping conflict with those of Monte et al. (Monte et al., 2021), who suggest that knee EMA is independent of step frequency (2.0 – 3.5 Hz). Our data may differ from those of Monte et al. due to a difference in methodology. We calculated joint-specific average EMA during the stance phase when joint moments exceeded 25% of their peak value, whereas Monte et al. separated stance into two phases and included EMA values obtained when GRF and center of pressure are small (near ground contact or toe off), which increases variability in EMA and may obscure changes that occur with step frequency (Griffin et al., 2003). There may have also been difference in inter-participant hopping strategies between studies, where participants may have kept their knees “locked” or “unlocked”. While our average knee EMA data suggest that participants straighten their legs to hop at faster step frequencies, three of our participants did not appreciably change their knee EMA across step frequency (Fig. 5). This may suggest that some of our participants choose a “locked” knee strategy and that the difference in participants and possibly strategies between the two studies could have been due to chance. Further research is warranted to determine if a difference in hopping strategy could explain the difference in knee EMA between studies.

The “cost of generating force” hypothesis originally put forth by Kram & Taylor (Kram and Taylor, 1990) provides a simple equation (Eqn. 1) that links biomechanics to metabolic energy expenditure across running velocities. The equation assumes that across running speeds, animals employ a constant EMA and muscles operate at similar shortening velocities. Each of these assumptions affects metabolic cost (Taylor, 1994), but do not detract from the elegance of a simple equation to well predict the metabolic cost of running in different sized animals across velocities. By addressing these assumptions, the accuracy for predicting metabolic energy expenditure for running and hopping at different step frequencies could be improved (Kipp et al., 2018b; Roberts et al., 1998b; Wright and Weyand, 2001).

The cost-coefficients, encompass factors that influence muscle metabolic energy expenditure per unit active muscle volume (J□cm^3^). As such, the values of these coefficients change when unaccounted factors that affect metabolic energy expenditure change. For example, changes in muscle force-length-velocity affect metabolic energy expenditure per unit of force production but are not accounted for in equations 1 & 2. In the present study, we show that accounting for changes in *V*_*m*_ (through EMA) and *t*_*c*_^*-1*^ (Eqn. 2) yields a constant cost-coefficient, *k*, across step-frequency (Fig. 7). Thus, by incorporating EMA into Eqn. 2, the metabolic energy expenditure per unit active muscle volume is consistent across step frequencies, unlike in the simpler Eqn. 1 (Fig. 7). It may be possible for future studies, using ultrasound or modeling approaches to account for additional assumptions, such as muscle fiber shortening velocity and length to further refine the “cost of generating force” hypothesis.

Changes in muscle contractile dynamics (i.e., muscle shortening velocity and average operating length) likely influence total active muscle volume. In a recent study, Beck et al (Beck et al., 2020) demonstrated that producing the same cycle-average forces with a decreasing duty factor (the product of contact time and frequency) during cyclic soleus contractions requires greater peak muscle force, a decrease in fascicle operating length, and a general increase in active muscle volume and metabolic energy expenditure. The authors proposed that accounting for duty factor may improve the calculation of active muscle volume by providing a surrogate for muscle contractile dynamics; thereby addressing one of the assumptions of the “cost of generating force” hypothesis.

A potential limitation of our study is the use of static internal muscle-tendon moment arms, fascicle lengths, and pennation angles to estimate active muscle volume. We intentionally did this to allow a direct comparison of our results to those of previous studies (Biewener et al., 2004; Kipp et al., 2018a) that account for active muscle volume changes during human locomotion. We found that accounting for active muscle volume increases inter-participant variability of predicted metabolic energy expenditure compared to assuming constant active muscle volume (Fig. 2 & 3). This increase in variability may be due to the assumption of fixed-length, muscle moment arms at each joint. Inter-participant variability in total active muscle volume and predicted metabolic energy expenditure using Eqn. 2 might be reduced by accounting for changes in muscle moment arms during the stance phase. Previous studies have shown that muscle-tendon moment arms change with joint angle (Arnold et al., 2010; Hoy et al., 1990; Rasske et al., 2017). Thus, using variable muscle-tendon moment arms that change with joint angle could further improve the estimate of active muscle volume and metabolic energy expenditure.

## Conclusion

In this study, we evaluated the “cost of generating force” hypothesis for predicting metabolic energy expenditure across different step frequencies during running and hopping. We found that accounting for changes in effective mechanical advantage to compute active muscle volume resulted in a near-constant cost-coefficient, *k*, and improved the estimation of participant metabolic energy expenditure across step frequencies.

## List of symbols and abbreviations

c: cost-coefficient
EMA: effective mechanical advantage
Ė_met_: metabolic power
F_BW_: force in units of body weight
F_mtu_: muscle-tendon force
GRF: ground reaction force
k: cost-coefficient
L: fascicle length
M: joint moment
PCSA: physiological cross-sectional area
PSF: preferred step frequency
r: muscle-tendon moment arm
R: GRF moment arm
t_c_: ground contact time
t_c_^-1^: rate of muscle force production
V_m_: active muscle volume
σ: muscle stress

## Author Contributions

Conceptualization: S.P.A., O.N.B., A.M.G; Methodology: S.P.A., O.N.B., A.M.G; Investigation: S.P.A., O.N.B; Formal analysis: S.P.A; Writing – original draft: S.P.A; Writing – review & editing: S.P.A., O.N.B., A.M.G; Visualization: S.P.A; Supervision: A.M.G.

